# High-throughput screening for Cushing’s disease: therapeutic potential of thiostrepton via cell cycle regulation

**DOI:** 10.1101/2024.02.22.581351

**Authors:** Takuro Hakata, Ichiro Yamauchi, Daisuke Kosugi, Taku Sugawa, Haruka Fujita, Kentaro Okamoto, Yohei Ueda, Toshihito Fujii, Daisuke Taura, Nobuya Inagaki

**Affiliations:** Department of Diabetes, Endocrinology and Nutrition, Kyoto University Graduate School of Medicine, 54 Kawaharacho, Syogoin, Sakyo-ku, Kyoto 606-8507, Japan; Medical Research Institute KITANO HOSPITAL, PIIF Tazuke-kofukai, 2-4-20 Ohgimachi, Kita-ku, Osaka 530-8480, Japan

**Keywords:** high-throughput screening, Cushing’s disease, thiostrepton, cell cycle, CDK inhibitor

## Abstract

Cushing’s disease is a life-threatening disorder caused by autonomous secretion of adrenocorticotropic hormone (ACTH) from pituitary neuroendocrine tumors (PitNETs). Few drugs are indicated for inoperative Cushing’s disease, in particular that due to aggressive PitNETs. To explore agents that regulate ACTH-secreting PitNETs, we conducted high-throughput screening (HTS) using AtT-20, a murine pituitary tumor cell line characterized by ACTH secretion. For the HTS, we constructed a live cell– based ACTH reporter assay for high-throughput evaluation of ACTH changes. This assay was based on HEK293T cells overexpressing components of the ACTH receptor and a fluorescent cAMP biosensor, with high-throughput acquisition of fluorescence images at the single-cell level. Of 2480 screened bioactive compounds, over 50% inhibition of ACTH secreted from AtT-20 cells was seen with 84 compounds at 10 μM, and 20 compounds at 1 μM. Among these hit compounds, we focused on thiostrepton (TS) and determined its antitumor effects in both *in vitro* and *in vivo* xenograft models of Cushing’s disease. Transcriptome and flow cytometry analyses revealed that TS administration induced AtT-20 cell cycle arrest at the G2/M phase, which was mediated by FOXM1-independent mechanisms including downregulation of cyclins. Simultaneous TS administration with a CDK 4/6 inhibitor that affected the cell cycle at the G0/1 phase showed cooperative antitumor effects. Thus, TS is a promising therapeutic agent for Cushing’s disease. Our list of hit compounds and new mechanistic insights into TS effects serve as a valuable foundation for future research.

## Introduction

Cushing’s disease results from excessive cortisol production by pituitary neuroendocrine tumors (PitNETs) that secrete adrenocorticotropic hormone (ACTH). Patients with Cushing’s disease exhibit various clinical features of chronic hypercortisolism, including central obesity, moon face, glucose intolerance, hypertension, dyslipidemia, osteoporosis, depression, and an immunocompromised state (1). If not treated, Cushing’s disease is associated with a high mortality rate, primarily due to cardiovascular disease or infections (2, 3). Transsphenoidal surgery is the first-line treatment recommended for Cushing’s disease (4). The remission rate is approximately 80% for micro PitNETs, and around 60% for macro PitNETs. Furthermore, repeat surgery is associated with less favorable outcomes (5). Thus, some patients with persistent or recurrent Cushing’s disease require additional treatment with medical therapies.

In the management of hypercortisolism due to Cushing’s disease, adrenal-targeting drugs such as metyrapone, ketoconazole, and osilodrostat are used for acute-phase intervention but do not directly affect ACTH-secreting PitNETs (4, 6). To achieve long-term regulation, drugs targeting ACTH-secreting PitNETs are necessary. Among such drugs, pasireotide, a somatostatin receptor ligand, has demonstrated efficacy in reducing ACTH production and tumor volume, with response rates ranging from 20%– 52% (7–10). Cabergoline, a dopamine receptor agonist, normalized urinary cortisol in 30–40% of patients (11–13). Both drugs are associated with adverse events, including severe hyperglycemia, diarrhea, and long QT syndrome with pasireotide, and nausea, asthenia, and vertigo with cabergoline (5). Moreover, these agents may have an insufficient effect on the aggressive or metastatic PitNETs that most often secrete ACTH (14). Temozolomide, an alkylating agent, is sometimes used for aggressive and metastatic PitNETs. Although a significant volume reduction of PitNETs was seen in approximately 40% of patients treated with temozolomide, complete tumor regression was achieved in only about 5% of patients (14, 15). Therefore, there is an unmet need for agents that can effectively treat Cushing’s disease caused by aggressive PitNETs.

High-throughput screening (HTS) is a powerful technique for identifying therapeutic targets. Here we aimed to perform HTS to discover novel agents that suppress ACTH secretion and inhibit the growth of ACTH-secreting PitNETs. To construct an HTS assay, we focused on the signaling mechanisms of the ACTH receptor. This receptor consists of two components, the melanocortin 2 receptor (MC2R) and the melanocortin 2 receptor accessory protein (MRAP), which cooperatively act as a G protein–coupled receptor (16). ACTH binding to the ACTH receptor causes increased intracellular levels of cyclic adenosine monophosphate (cAMP), a second messenger. In this context, we constructed an ACTH reporter assay based on evaluating cAMP produced by stimulation of the ACTH receptor, as described in the following sections.

In the present study, we performed HTS based on our ACTH reporter assay using AtT-20, a murine pituitary tumor cell line characterized by ACTH secretion. As a candidate agent to treat Cushing’s disease, we identified thiostrepton (TS) from among 2480 bioactive compounds. TS exhibited significant antitumor effects in both *in vitro* and *in vivo* models of Cushing’s disease. Furthermore, we elucidated the molecular mechanisms of TS effects, which include downregulation of cyclins.

## Results

### Construction of the ACTH reporter assay

As stated in the Introduction, we constructed an ACTH reporter assay for high-throughput measurement of ACTH concentrations. This is a live cell–based assay that measured fluctuations in cAMP levels caused by stimulation of the ACTH receptor. Detailed information on the ACTH reporter assay is presented in Figure 1A. We transfected expression vectors containing the ACTH receptor and a cAMP biosensor into HEK293T cells. MC2R and MRAP, components of the ACTH receptor, were expressed by transfecting the respective vectors. Pink Flamindo was used as the cAMP biosensor (17). Stimulation of the transfected cells by ACTH resulted in increased intracellular cAMP levels, which then augmented the fluorescence of the cAMP biosensor. High-throughput acquisition of images at the single-cell level was performed using an Opera Phenix High-Content Screening System, and changes in the fluorescence intensity of individual cells were compared before and after stimulation (See also Materials and Methods).

**Figure 1.**
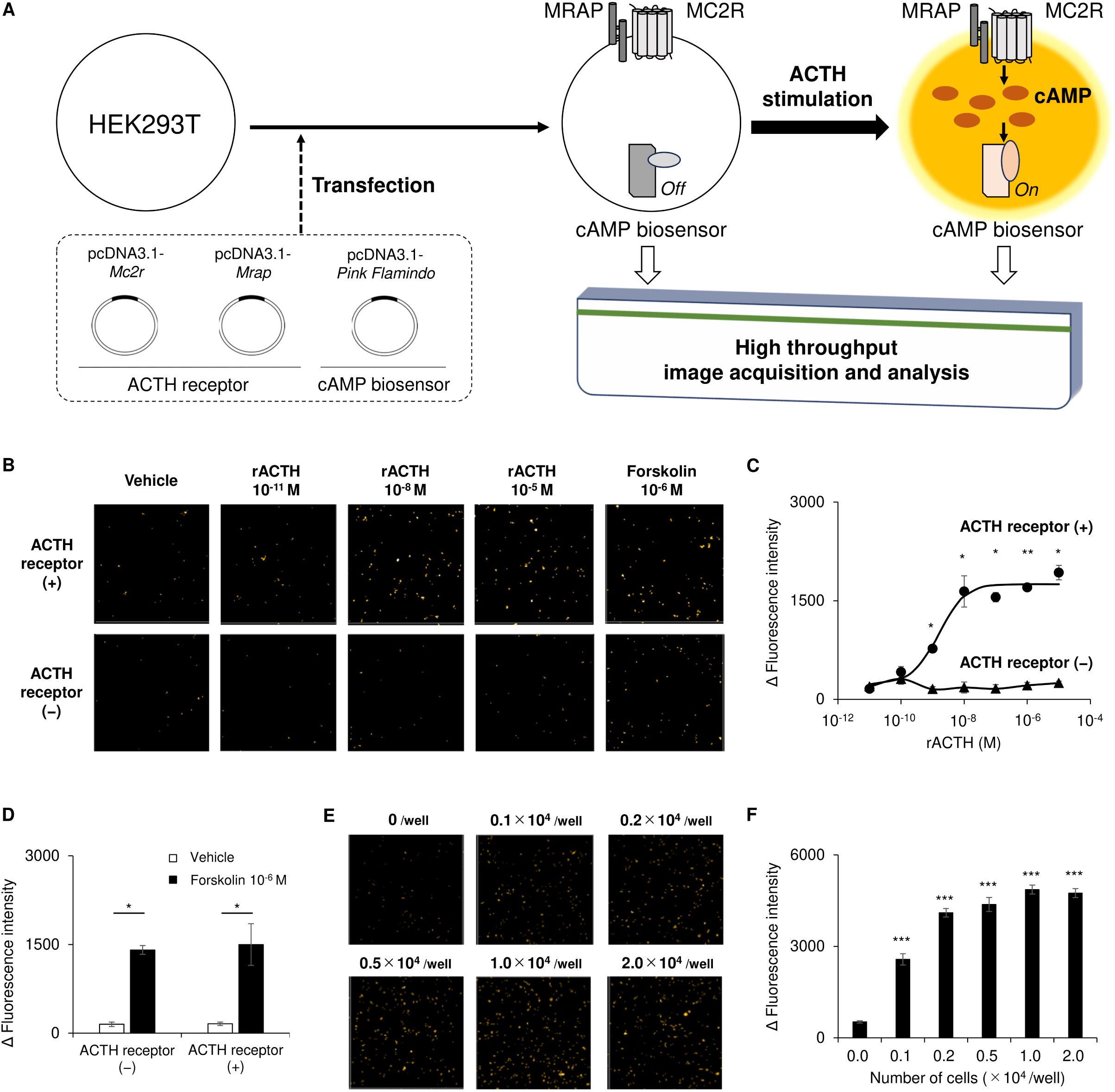
Validation of the ACTH reporter assay constructed for high-throughput screening (HTS). (A) An assay flow diagram of the ACTH reporter assay. ACTH, adrenocorticotropic hormone; MC2R, melanocortin 2 receptor; MRAP, melanocortin 2 receptor accessory protein; and cAMP, cyclic adenosine monophosphate. (B) Images of fluorescence by the cAMP biosensor in HEK293T cells transfected with the ACTH receptor (ACTH receptor (+)) and those with an empty vector (ACTH receptor (−)). A diluted series of recombinant ACTH (rACTH) or forskolin was administered to the cells. (C) Sigmoidal curves derived from changes in fluorescence intensity (Δ fluorescence intensity) of the transfected HEK293T cells after administration of rACTH. *n* = 3 each. (D) Δ fluorescence intensity of the transfected HEK293T cells after administration of forskolin. *n* = 3 each. (E) Fluorescence images of ACTH receptor (+) HEK293T cells treated with conditioned media from various numbers of AtT-20 cells. (F) Δ fluorescence intensity of ACTH receptor (+) HEK293T cells treated with conditioned media from AtT-20 cells. *n* = 6 each. Data are represented as means ± SEM. Statistical analyses were performed using the Student’s *t*-test for panels C and D and ANOVA followed by the Dunnett test with comparisons to conditioned media without AtT-20 cells for panel F. The *p*-value is presented as \**p* < 0.05, \*\**p* < 0.01, and \*\*\**p* < 0.001.

We validated this ACTH reporter assay by comparing two types of HEK293T cells: those transfected with both the cAMP biosensor and the ACTH receptor, and those transfected with the cAMP biosensor only (Figure 1B–1D). Administration of recombinant ACTH significantly and dose-dependently augmented the fluorescence intensity of HEK293T cells transfected with the ACTH receptor, whereas HEK293T cells without the ACTH receptor did not respond (Figure 1B, 1C). In the presence of forskolin, augmentation of fluorescence intensity was observed in both types of cells, indicating the functionality of the cAMP biosensor (Figure 1D). In addition, to implement HTS, we used the ACTH reporter assay to evaluate ACTH concentrations in conditioned media from AtT-20 cells. As the number of AtT-20 cells per well increased, their conditioned media augmented the fluorescence intensity of HEK293T cells transfected with the ACTH reporter to a plateau of 1.0 × 10^4^ cells/well (Figure 1E, 1F). Thus, we confirmed that the ACTH reporter assay was applicable to HTS using AtT-20 cells, and determined that the optimal number of AtT-20 cells used in HTS was 1.0 × 10^4^ cells/well.

### HTS using AtT-20 cells

The HTS process is outlined in Figure 2A. AtT-20 cells were exposed to each compound and incubated for 24 h. Following incubation, ACTH concentrations in the conditioned media from the cells were evaluated using the ACTH reporter assay. The first screening of all 2480 compounds in the chemical library at 10 μM was conducted in duplicate. Of the 2480 screened compounds, 254 exhibited more than 50% inhibition of ACTH secretion (Figure 2B). We calculated the Z’ factor, a statistical parameter reflecting assay performance, based on the results obtained using positive and negative control media (Supplementary Figure 1A): the positive control consisted of conditioned media from AtT-20 cells treated with vehicle, while the negative control comprised media incubated without AtT-20 cells. The mean Z’ factor of all plates was 0.50 ± 0.03. We concurrently evaluated cell viability and found that 213 compounds resulted in more than 50% inhibition of cell viability (Supplementary Figure 1B). Interestingly, changes in ACTH secretion strongly correlated with cell viability (r = 0.92, *p* < 0.001) (Supplementary Figure 1C).

**Figure 2.**
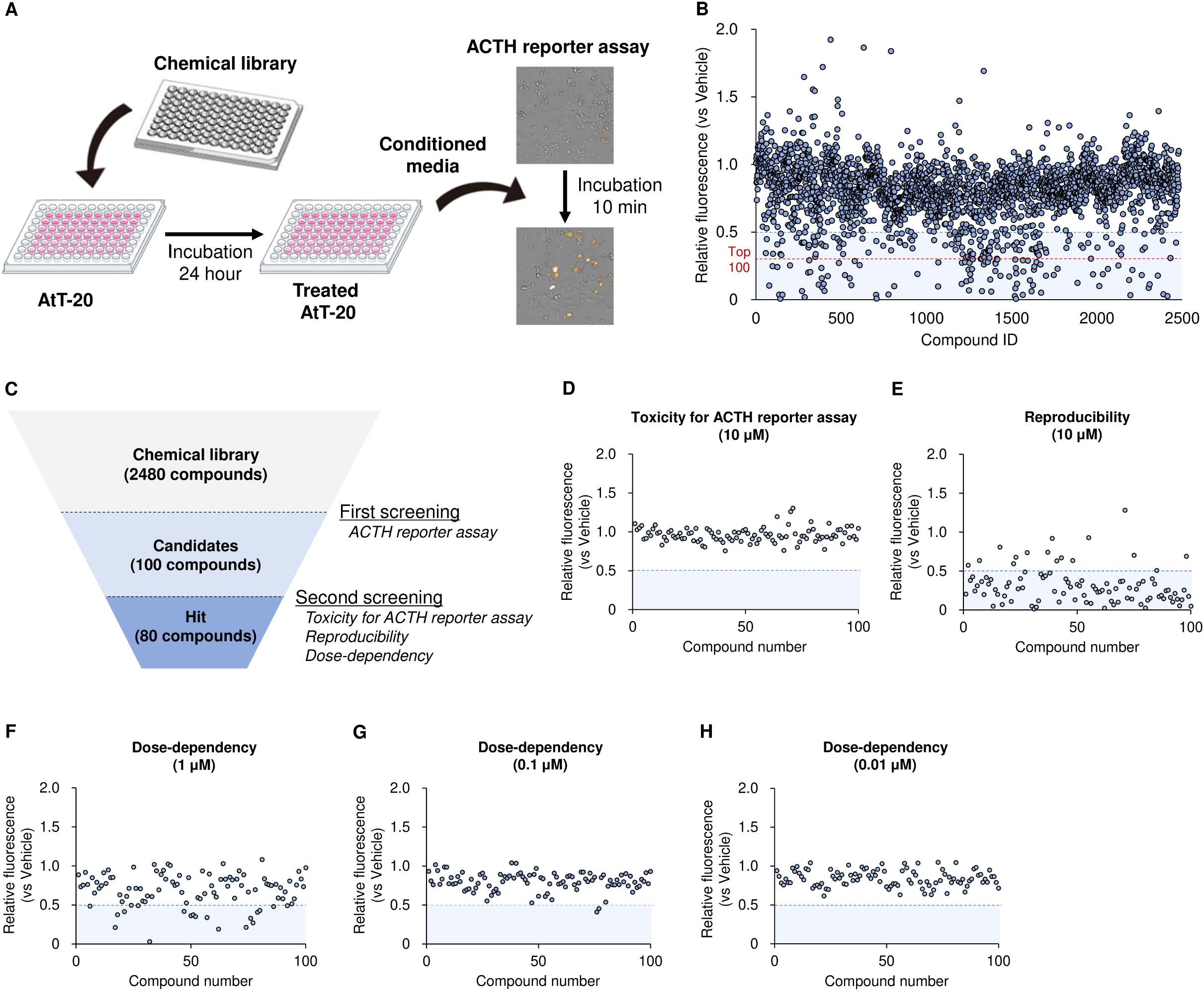
Results of HTS of AtT-20 cells. (A) An assay flow diagram of HTS. (B) Initial screening for all 2480 compounds in the chemical library. Values were normalized as relative fluorescence; in each plate, 1.0 was the result of AtT-20 cells treated with vehicle. The blue dotted line indicates 0.5 x relative fluorescence, which we considered to indicate a significant reduction. The red dotted line indicates the cut-off for the top 100 compounds in terms of inhibiting ACTH secretion. (C) A schema of HTS results including the second screening whose results are shown in panels D–H. (D) Evaluation of direct toxicity in the ACTH reporter assay. The top 100 compounds were administered at 10 μM with 10^-8^ M of recombinant ACTH. (E) Verification of reproducibility using the HTS assay. The top 100 compounds were administered at 10 μM. (F–H) Evaluation of dose dependency in the HTS assay. The top 100 compounds were administered at 1 μM (F), 0.1 μM (G), and 0.01 μM (H). Assays were performed in duplicate and results were calculated from their means.

We obtained a number of candidate compounds that might inhibit ACTH secretion by PitNETs, and then performed a second screening for the top 100 compounds (Figure 2C). First, we examined whether compounds carried over to the conditioned media displayed toxicity against the live cells used in the ACTH reporter assay. We performed the ACTH reporter assay using both recombinant ACTH and the candidate compounds, and the results did not suggest significant toxicity of any of the top 100 compounds examined (Figure 2D). Second, we validated the reproducibility of the first screening by again administering the compounds at 10 μM, and found that 84 of the 100 compounds consistently exhibited more than 50% inhibition of ACTH secretion (Figure 2E). Third, we examined dose dependency and observed that 20 compounds resulted in more than 50% inhibition of ACTH at 1 μM, 2 compounds at 0.1 μM, and no compounds at 0.01 μM (Figure 2F–2H).

Thus, we identified 84 hit compounds that repeatedly exhibited more than 50% inhibition of ACTH secretion (Supplementary Table 1). In particular, 20 hit compounds with significant inhibitory effects at 1 μM are listed in Table 1.

**Table 1.** Hit compounds administered at a concentration of 1 μM.

| **Compound** | **Classification** | **ACTH**<br>**inhibition** | **Cell viability**<br>**inhibition** | **IC50**<br>**(μM)** |
| --- | --- | --- | --- | --- |
| Belinostat (PXD101) | HDAC inhibitor | 73% | 70% | 0.50 |
| Geldanamycin | HSP90 inhibitor | 95% | 83% | 0.23 |
| GSK2126458 | PI3K and mTOR inhibitor | 72% | 77% | 0.03 |
| PF-05212384 (PKI-587) | PI3K and mTOR inhibitor | 76% | 69% | 0.05 |
| GDC-0980 (RG7422) | PI3K and mTOR inhibitor | 74% | 76% | 0.64 |
| Lacidipine | Ca channel blocker | 72% | 19% | 0.77 |
| Mibefradil dihydrochloride hydrate | Ca channel blocker | 97% | 79% | 0.90 |
| GBR 12909 dihydrochloride | Dopamine reuptake inhibitor | 84% | 69% | 0.41 |
| ON-01910 | PLK inhibitor | 86% | 78% | 0.18 |
| Camptothecine | Cytotoxic anticancer drug | 74% | 79% | 0.28 |
| Mitoxantrone | Cytotoxic anticancer drug | 98% | 49% | 0.29 |
| Mitoxantrone dihydrochloride | Cytotoxic anticancer drug | 66% | 100% | 0.24 |
| Vincristine | Cytotoxic anticancer drug | 72% | 76% | 0.55 |
| MG-132 | Proteasome inhibitor | 89% | 69% | 0.34 |
| Bortezomib | Proteasome inhibitor | 79% | 78% | 0.53 |
| Thiostrepton | Antibiotics | 82% | 58% | 0.99 |
| Haloprogin | Antifungal drug | 94% | 72% | 0.45 |
| Auranofin | Antirheumatic drug | 90% | 78% | 0.18 |
| Chlorhexidine | Disinfectant | 99% | 73% | 0.89 |
| Plumbagin | Toxic substance | 81% | 84% | 0.86 |

IC50, 50% inhibitory concentration; HDAC, histone deacetylase; HSP90, heat shock protein 90; PI3K, phosphatidylinositol-3 kinase; mTOR, mammalian target of rapamycin; PLK, polo-like kinase.

### Verification of TS effects on AtT-20 cells

Among the 20 compounds with significant effects at 1 μM, histone deacetylase (HDAC) inhibitors (18–21), heat shock protein 90 (HSP90) inhibitors (22–24), phosphatidylinositol-3 kinase (PI3K) inhibitors (18), and proteasome inhibitors (25) have been reported to have antitumor effects against ACTH-secreting PitNETs. After excluding the compounds belonging to these classes, we focused on TS, a natural compound containing a thiazole ring. TS was isolated from *Streptomyces* species and was approved by the Food and Drug Administration as an antibiotic in animals. Notably, TS has shown antitumor effects in various cancers, including breast cancer (26, 27), ovarian cancer (28, 29), and meningioma (30). In this context, we aimed to elucidate the potential of TS to treat ACTH-secreting PitNETs.

We re-administered TS to AtT-20 cells to confirm its effects *in vitro*. We used the ACTH reporter assay to analyze its ability to dose-dependently inhibit ACTH secretion, and determined that the 50% inhibitory concentration (IC50) was 0.9 μM (Figure 3A). Measurement of secreted ACTH in conditioned media by conventional ELISA yielded a similar IC50 of 1.3 μM (Figure 3B). However, TS did not decrease the expression of *Pomc*, which encodes the ACTH precursor (Figure 3C). We considered that TS inhibited ACTH secretion via antitumor effects and analyzed changes in cell number and viability. The number of AtT-20 cells was decreased by TS administration in a dose-dependent manner after both 24 and 48 h (Supplementary Figure 2A, 2B). Since cell viability also decreased dose dependently at both time points (Figure 3D), we proceeded following investigation by evaluating cell viability.

**Figure 3.**
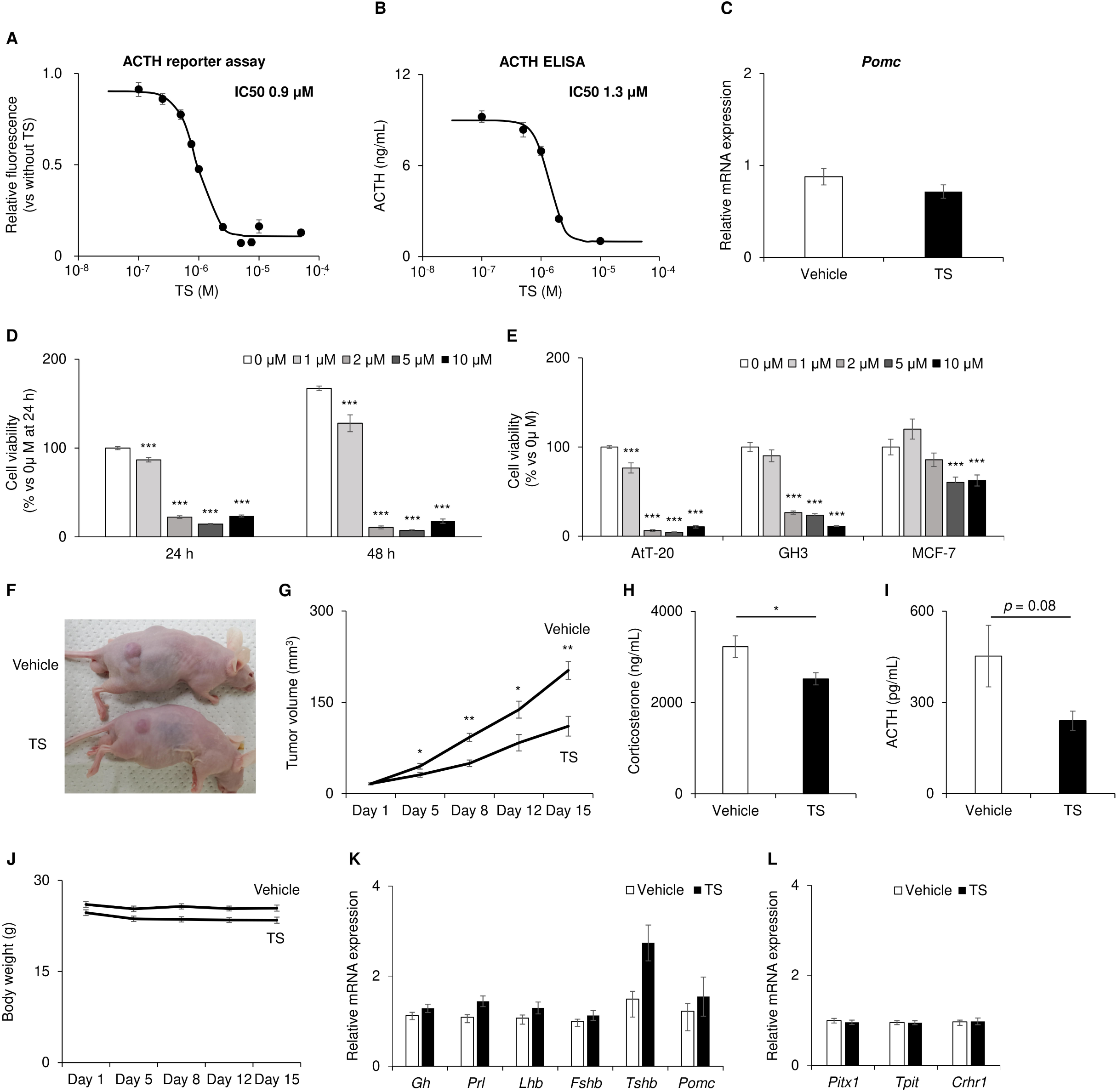
Effects of thiostrepton (TS) in *in vitro* and *in vivo* models of Cushing’s disease. (A–E) Elucidation of TS effects on AtT-20 cells *in vitro*. (A, B) Evaluation of changes in ACTH secreted by AtT-20 cells after treatment with a dilution series with TS for 24 h. The 50% inhibitory concentration (IC50) was calculated using sigmoidal curves. An ACTH reporter assay (A) and ELISA (B) were performed. Results in panel A are shown as relative fluorescence: 1.0 was set as the value without TS administration. *n* = 8 for each group in panel A, and *n* = 4 for each group in panel B. (C) Quantitative RT-PCR for *Pomc* mRNA of AtT-20 cells treated with 2 μM TS for 24 h. *n* = 6 each. (D) Cell viability of AtT-20 cells treated with a dilution series of TS at 24 and 48 h. Results are shown as the ratio relative to 0 μM at 24 h. *n* = 8 each. (E) Comparisons of TS effects on the viability of AtT-20, GH3, and MCF-7 cells. Cells were treated with a dilution series of TS for 48 h. Results are shown as the ratio relative to 0 μM in each cell. *n* = 8 each. (F–K) Verification of TS effects in an *in vivo* xenografted model. (F) A representative image of treated mice. Vehicle and 300 mg/kg of TS were administered every other day through intraperitoneal injection. *n* = 5 for vehicle group and *n* = 7 for TS group. (G) Growth curves of tumor volume. (H, I) Plasma levels of hormones in treated mice; corticosterone (H) and ACTH (I). (J) Growth curves of body weight. (K, L) Quantitative RT-PCR of mRNA expression in the pituitary glands of treated mice. mRNA levels of genes related to anterior pituitary hormones (K) and those of genes specific for ACTH-secreting cells (L). Data are represented as fold-change versus values of mice treated with vehicle. Data of quantitative RT-PCR are normalized by means of *Actb* and *Gapdh*. Data are represented as means ± SEM. Statistical analyses were performed using the Student’s *t*-test for panels C and G–L, and ANOVA followed by the Dunnett test with comparisons to results without TS administration for panels D and E. The *p*-value is presented as \**p* < 0.05, \*\**p* < 0.01, and \*\*\**p* < 0.001.

Furthermore, we administered TS to GH3, a rat pituitary cell line characterized by the secretion of growth hormone, and to MCF-7, a breast cancer cell line that was previously shown to be susceptible to TS (26). The IC50 after TS administration for 48 h was 1.0 μM in AtT-20 cells, 1.4 μM in GH3 cells, and not reached in MCF-7 cells (Figure 3E). We further determined changes in cell numbers after TS administration (Supplementary Figure 2A–2F): at 24 h, the IC50 was 1.4 μM in AtT-20 cells, 0.7 μM in GH3 cells, and not reached in MCF-7 cells, while at 48 h it was 1.1 μM in AtT-20 cells, 1.1 μM in GH3 cells, and not reached in MCF-7 cells. Thus, TS more strongly reduced the viability of AtT-20 and GH3 cells than that of MCF-7 cells.

Following verification *in vitro*, we conducted experiments to determine TS effects *in vivo*. We generated a tumor xenograft model by subcutaneous inoculation of AtT-20 cells into KSN/Slc mice (Figure 3F). TS administration significantly reduced tumor volume: 202.4 ± 14.7 mm^3^ with vehicle, 110.6 ± 14.2 mm^3^ with TS, *p* = 0.005 (Figure 3G). Weights of resected tumors were significantly lower with TS administration (91.4± 19.6 mg) than with vehicle (168.0 ± 25.2 mg) (*p* = 0.025). In addition, plasma corticosterone concentrations were significantly reduced by TS administration (Figure 3H): 3223.9 ± 239.4 pg/mL with vehicle, 2518.2 ± 132.4 pg/mL with TS, *p* = 0.031. Although it was possible that decreased corticosterone concentrations could increase ACTH secretion via a feedback loop, plasma ACTH concentrations were reduced by TS administration, though not significantly (Figure 3I); 452.1 ± 102.0 pg/mL with vehicle, 239.7 ± 31.6 pg/mL with TS, *p* = 0.08.

TS was not associated with checked toxic effects. Specifically, its administration did not significantly change body weights throughout the study period (Figure 3J), or the gene expressions of anterior pituitary hormones (Figure 3K). We further evaluated the expressions of genes specific to ACTH-secreting cells, and identified no changes in the mRNA levels of *Pitx1*, *Tbx19*, or *Crhr1* (Figure 3L).

Altogether, these results confirm the suppressive effects of TS on the growth of ACTH-secreting PitNETs in both *in vitro* and *in vivo* models of Cushing’s disease.

### Exploration of the mechanisms of the antitumor effects of TS

Subsequently, we explored the mechanisms of the antitumor effects of TS. Previous studies regarding various cancers reported that the primary effect of TS was the inhibition of FOXM1 (26–30). In AtT-20 cells, TS caused significant decreases in the expressions of both *Foxm1* mRNA and FOXM1 protein (Figure 4A, Supplementary Figure 3A, 3B). To clarify the function of FOXM1 in the growth of AtT-20 cells, we performed knockdown of *Foxm1* using short interfering RNAs (siRNAs). The efficacy of knockdown was validated based on the decreases in the expressions of *Foxm1* mRNA and FOXM1 protein (Figure 4B, Supplementary Figure 3C, 3D). *Foxm1* knockdown did not attenuate the viability of AtT-20 cells at 24 h, but did so at 48 h (Figure 4C). However, the magnitude to which *Foxm1* knockdown decreased cell viability appeared to be less pronounced compared to TS (Figure 3C).

**Figure 4.**
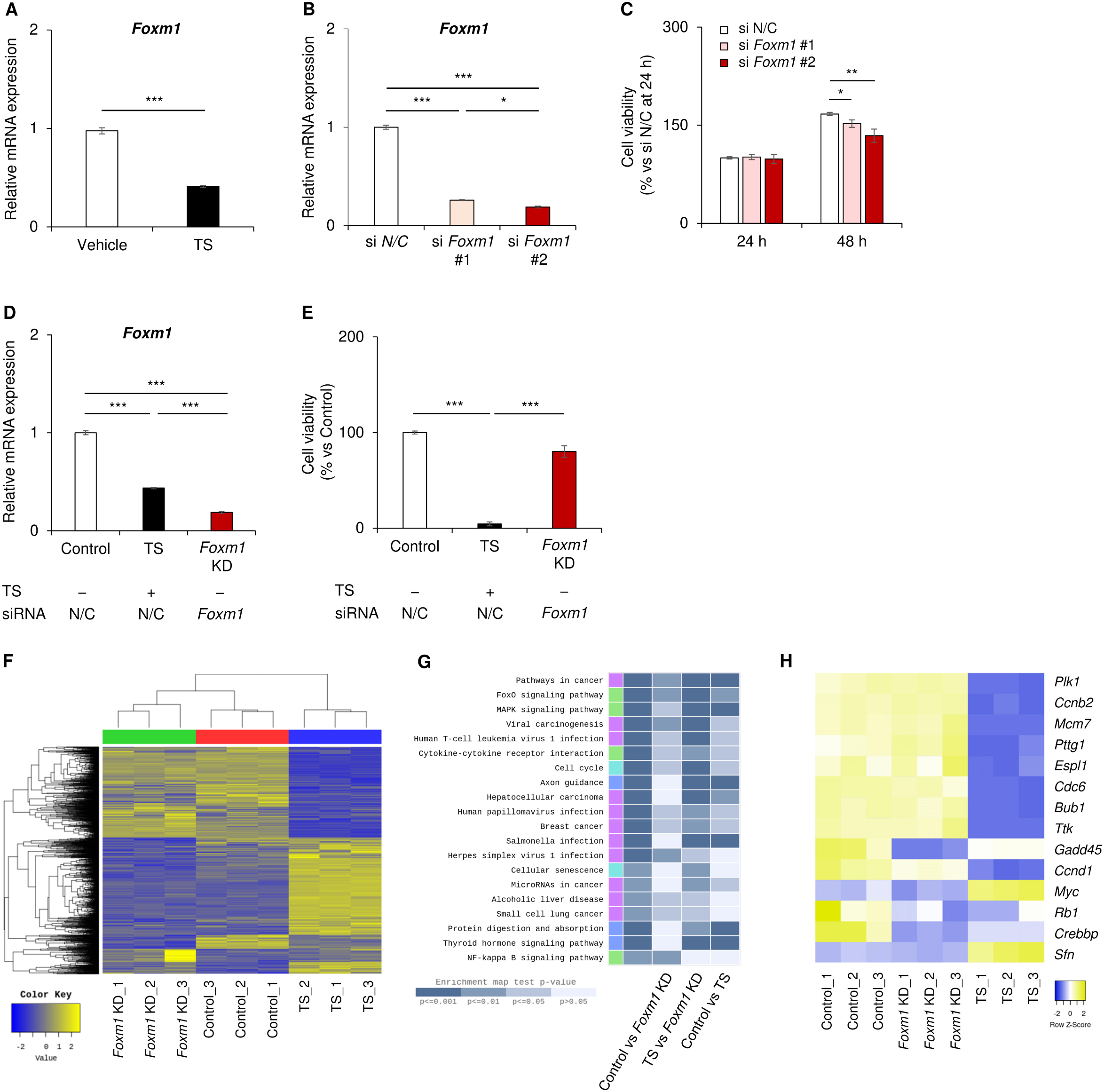
Exploring mechanisms of TS effects against AtT-20 cells. (A) Quantitative RT-PCR of *Foxm1* mRNA in AtT-20 cells treated with 2 μM TS for 24 h. *n* = 6 each. (B, C) Quantitative RT-PCR of *Foxm1* mRNA (B) and cell viability (C) of AtT-20 cells with *Foxm1* knockdown (KD) by transfection with short interfering RNA (siRNA). A negative control siRNA (si N/C) was used, along with two siRNAs for *Foxm1* (si *Foxm1* #1 and si *Foxm1* #2). RNA samples were collected after 24 h and cell viability was evaluated at both 24 and 48 h after transfections. Results are shown as the ratio relative to si N/C at 24 h. *n* = 6 each. (D, E) Quantitative RT-PCR of *Foxm1* mRNA (D) and cell viability (E) of AtT-20 cells treated with TS and/or with *Foxm1* knockdown. The Control group was treated with vehicle and transfected with si N/C, the TS group with 2 μM TS and si N/C, and the *Foxm1* KD group with vehicle and si *Foxm1* #2. *n* = 6 each. (F–H) Results of RNA-sequencing (RNA-seq) using RNA samples obtained in the experiments shown in panels D and E. Samples were analyzed as a mix of every two samples. *n* = 3 each. Panels F–H show hierarchical clustering (F), the top 20 terms from the enrichment analysis by the Kyoto Encyclopedia of Genes and Genomes (KEGG) (G), and DEGs associated with the cell cycle pathway in KEGG (H). DEGs were determined as an adjusted *p* < 0.05 and fold changes ≤ −2 or ≥ 2. Data of quantitative RT-PCR are normalized by means of *Actb* and *Gapdh*. Data are represented as means ± SEM. Statistical analyses were performed using the Student’s *t*-test for a panel A, and ANOVA followed by the Tukey-Kramer test for panels B–E. The *p*-value is presented as \**p* < 0.05, \*\**p* < 0.01, and \*\*\**p* < 0.001.

Therefore, we proceeded to directly compare the effects of TS administration (TS group) and *Foxm1* knockdown (*Foxm1* KD group). We prepared the Control group without either TS administration or *Foxm1* knockdown. In this experiment, siRNA #2 for *Foxm1* was used because it provided more significant *Foxm1* knockdown (Supplementary Figure 3C, 3D) and less cell viability (Figure 4C) than siRNA #1. Both TS administration and *Foxm1* knockdown significantly decreased *Foxm1* mRNA. *Foxm1* knockdown rather strongly decreased *Foxm1* mRNA compared to TS administration (Figure 4D). On the other hand, cell viability was not significantly reduced by *Foxm1* knockdown, but was significantly reduced by TS administration (Figure 4E). Based on these results, we hypothesized that TS predominantly attenuates cell viability by targeting molecules other than FOXM1.

We performed transcriptome analysis using RNA-sequencing (RNA-seq) to elucidate additional pathways affected by TS. We provide the RNA-seq dataset in Supplementary Table 2. Hierarchical clustering analysis revealed clear differences between the Control and TS groups, while fewer differences were observed between the Control and *Foxm1* KD groups (Figure 4F). Regarding differentially expressed genes (DEGs), the TS group contained 521 upregulated genes and 496 downregulated genes, whereas the *Foxm1* KD group contained only 100 upregulated genes and 157 downregulated genes (Supplementary Figure 4A–4C). Thus, as with decreases in cell viability, transcriptome alteration was more pronounced in the TS group compared with the *Foxm1* KD group.

Enrichment analyses were performed using the Kyoto Encyclopedia of Genes and Genomes (KEGG) and gene ontology (GO), and the top 20 terms are presented in Figure 4G and Supplementary Figure 4D–4F, respectively. Among these terms, we focused on cell cycle, which has been implicated in the pathogenesis of ACTH-secreting PitNETs in previous reports (31–33). DEGs associated with the cell cycle pathway are shown in Figure 4H. It was noteworthy that TS administration transcriptionally downregulated genes such as *Ccnb2* (34), *Ccnd1* (35), and *Pttg1* (36), which are abundantly expressed in aggressive PitNETs and related to PitNET tumorigenesis.

### TS effects on cell cycle in AtT-20 cells

Similar to our observation showing the downregulation of gene expressions of some cyclins (Figure 4H), previous reports demonstrated the upregulation of cyclins in ACTH-secreting PitNETs (31, 33, 35). We reviewed the expression levels of cyclin genes in RNA-seq analyses and found that almost all cyclins were downregulated in response to TS administration, while this downregulation was not observed with *Foxm1* knockdown (Figure 5A). Quantitative RT-PCR analyses confirmed that mRNA levels of *Ccna2*, *Ccnb2*, *Ccnd1*, and *Ccne1* were decreased by TS administration regardless of FOXM1 expression (Figure 5B).

**Figure 5.**
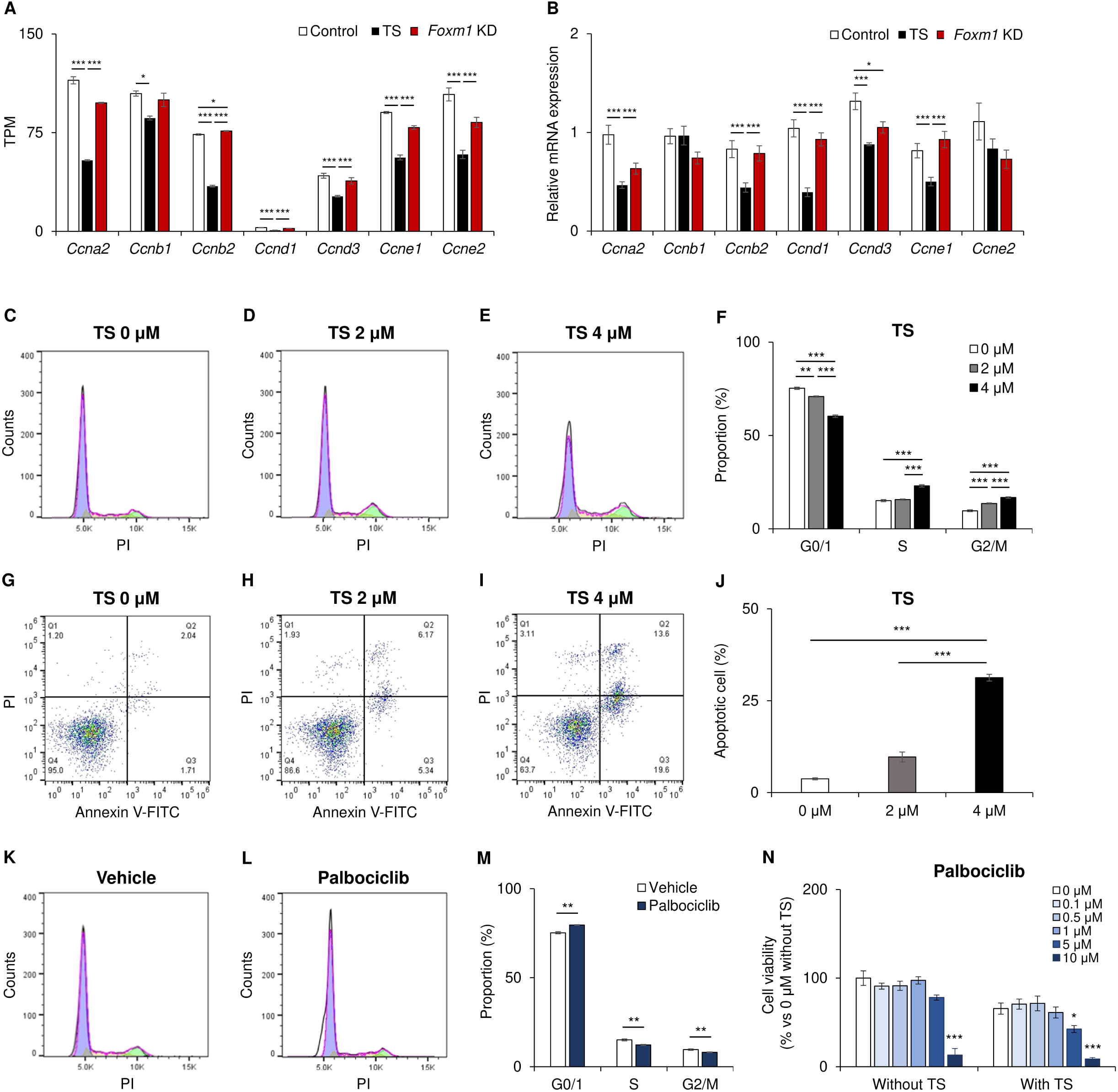
TS effects on the AtT-20 cell cycle. (A, B) Gene expression levels of cyclins in the Control, TS, and *Foxm1* KD groups. Data are represented as transcript per kilobase million (TPM) for RNA-seq (A), and as relative expression levels compared to the Control group for quantitative RT-PCR (B). *n* = 3 for each group in panel A, and *n* = 6 for each group in panel B. (C–F) Cell cycle analyses using propidium iodide (PI) staining and flow cytometry for AtT-20 cells treated with TS (0, 2, and 4 μM) for 24 h. *n* = 6 each. (G–J) Evaluation of apoptosis using staining with Annexin V-FITC and PI and flow cytometry for AtT-20 cells treated with TS (0, 2, and 4 μM) for 48 h. *n* = 3 each. (K–M) Cell cycle analyses using PI staining and flow cytometry for AtT-20 cells treated with 5 μM palbociclib for 24 h. *n* = 6 each. (N) Cell viability of AtT-20 cells treated with a dilution series of palbociclib. Either vehicle or 2 μM TS was co-administered with palbociclib and analysis was performed after 48 h. Results are shown as the ratio relative to values without palbociclib and TS. *n* = 6 each. Data of quantitative RT-PCR are normalized by means of *Actb* and *Gapdh*. Data are represented as means ± SEM. Statistical analyses were performed using ANOVA followed by the Tukey-Kramer test for panels A, B, F, and J, the Student’s *t*-test for panel M, and ANOVA followed by the Dunnett test with comparisons to results without palbociclib administration for panel N. The *p*-value is presented as \**p* < 0.05, \*\**p* < 0.01, and \*\*\**p* < 0.001.

To clarify the role of cyclin inhibition in TS effects, we analyzed cell cycle progression and apoptosis. TS administration altered the cell cycle distribution, specifically by dose-dependently increasing the proportion of cells in the G2/M and S phases and decreasing that of cells in the G0/1 phase (Figure 5C–5F). Meanwhile, Annexin V staining assays revealed that TS administration dose-dependently increased the proportion of apoptotic cells (Figure 5G–5J). Taken together, we considered that TS could induce cell cycle arrest in AtT-20 cells at the G2/M phase, and then induce apoptosis by decreasing cyclins via a FOXM1-independent pathway.

We next examined the interaction of TS with cyclin-dependent kinase (CDK) inhibitors, which are known to regulate the cell cycle and inhibit the growth of tumors caused by AtT-20 cells (27, 32, 33, 37). We specifically focused on palbociclib, an inhibitor of CDK4/6 that is clinically used in breast cancer (38). Palbociclib predominantly exerts its antitumor effects during the G0/1 phase (39, 40). We expected that co-administration of TS would enhance the effects of palbociclib because TS induces cell cycle arrest through another action in the G2/M phase. Cell cycle analyses in AtT-20 cells revealed that palbociclib increased the proportion of cells in the G0/1 phase and decreased that of cells in the S and G2/M phases (Figure 5K–5M). In addition, gene expressions of cyclins were not changed by palbociclib administration (Supplementary Figure 5A). Thereafter, we administered palbociclib to AtT-20 cells with or without TS. Palbociclib administration without TS reduced cell viability only at a concentration of 10 μM (Figure 5N). Notably, when TS was co-administered, palbociclib administration significantly reduced cell viability even at 5 μM (Figure 5N). Such cooperative inhibitory effects when co-administered with TS were not observed with other CDK inhibitors targeting the G2/M phase, the CDK 2/5 inhibitor seliciclib (R-roscovitine) (Supplementary Figure 5B), or the multiple CDK inhibitor flavopiridol (Supplementary Figure 5C).

## Discussion

Here we performed HTS to explore novel therapeutic agents for Cushing’s disease. Our HTS was conducted against AtT-20 cells as a model of ACTH-secreting PitNET. To complete HTS of 2480 chemical compounds, we established an ACTH reporter assay to evaluate ACTH concentrations in a high-throughput manner. Among the hit compounds, we focused on TS, an antimicrobial agent. TS exhibited significant therapeutic effects in both *in vitro* and *in vivo* models of Cushing’s disease. Transcriptome and flow cytometry analyses revealed that TS downregulated cyclins and then induced cell cycle arrest at the G2/M phase.

The ACTH reporter assay established in the present study is a unique assay that uses a fluorescence biosensor. Considering that cAMP plays a crucial role in ACTH receptor signaling as an intracellular second messenger, we evaluated cAMP elevation as a surrogate indicator for ACTH receptor stimulation. To secure specificity for the ACTH receptor, we overexpressed its essential components, MC2R and MRAP, in HEK293T cells lacking the ACTH receptor. In our assay, increased cAMP levels resulting from ACTH stimulation were dependent on both MC2R and MRAP, and a previous study showed no significant response in cells overexpressing only one of them (16). To overcome a limitation that the number of cells overexpressing both MC2R and MRAP through transfection can vary, we used the Opera Phenix High-Content Screening System, which is a powerful HTS system that enabled us to quantify various morphological features (41). In our ACTH reporter assay, we identified transfected cells exhibiting biosensor fluorescence and restricted our analysis to them. In other words, our method could evaluate cells with successful transfections and reduced biases caused by transient transfections. In fact, we completed HTS with high Z’ factors categorized as an excellent assay (42).

There has been only one previous report on HTS for Cushing’s disease. Its authors screened a kinase inhibitor library consisting of 430 compounds (18). Our screening included 2480 bioactive compounds, covering a broader range of drugs, thus allowing us to explore various classes (43). Some of the hit compounds listed in Table 1 and Supplementary Table 1 have been validated in terms of their potent efficacy against ACTH-secreting PitNETs; these include HDAC inhibitors (18–21), HSP90 inhibitors (22–24), PI3K inhibitors (18), proteasome inhibitors (25), an adrenergic receptor antagonist (44), and epidermal growth factor receptor inhibitors (45, 46). These results support the relevancy of our HTS results in terms of identifying therapeutic agents for Cushing’s disease.

Referring to our HTS results, we performed further examinations focusing on TS. TS administration attenuated the viability of AtT-20 cells and reduced ACTH secretion, but did not decrease *Pomc* mRNA levels. We considered that the inhibitory effects of TS on ACTH secretion seemed to be due to a decrease in the number of viable cells, and indeed, our overall HTS results showed that cell viability and ACTH secretion were positively correlated (Supplementary Figure 1C). TS administration was also effective *in vivo*, as determined using a xenograft model involving AtT-20 cells. Since specificity against tumor cells will be important when applying TS therapeutically, we verified that there were no changes in mRNA levels of pituitary hormones other than ACTH or those of transcription factors related to POMC expression. Moreover, mice administered TS did not exhibit apparent signs of toxicity such as loss of body weight.

The hypothesis that TS does not cause harmful effects on the normal pituitary gland is supported by several additional points, as follows. One mechanism of TS was downregulation of cyclins; this is considered to have minimal effects on normal pituitary tissue due to the fact that cyclin expression is lower in normal pituitary cells than in PitNETs (31, 33, 35). Furthermore, TS effects on AtT-20 cells were obtained at lower concentrations than with MCF-7 cells, in which TS effectiveness is well determined (26, 47). We believe that TS is a promising agent because its efficacy against AtT-20 cells was observed at relatively low concentrations and its toxicity was not severe at least. Interestingly, TS also reduced the cell viability of GH3, another PitNET cell line, at relatively low concentrations similar to those seen with AtT-20 cells. Cyclin genes have been shown to be upregulated not only in ACTH-secreting PitNETs but also in other types of PitNETs (35, 48–50). These results suggest that TS may have antitumor effects against various types of PitNETs.

As for the mechanisms of TS effects, FOXM1 was reported to be its predominant target (26–30). FOXM1 is a potential target in cancer treatment because it is highly expressed in some cancers, a finding associated with poor prognosis (51–58). FOXM1 plays a crucial role in maintaining cell homeostasis by regulating various biological processes, including proliferation, cell cycle progression, differentiation, DNA damage repair, and angiogenesis (59–61). FOXM1 is involved in cell entry to both the G1/S and G2/M phases by activating genes regulating these checkpoints (62–65). However, our direct comparisons of the TS and *Foxm1* KD groups revealed that FOXM1 contributed minimally to the antitumor effects of TS on AtT-20 cells.

We co-administered TS with CDK inhibitors to investigate the significance of cyclins in the effects of TS effect, as CDK acts with cyclins by forming various complexes. Palbociclib is a clinically used CDK4/6 inhibitor that primarily affects the G0/1 phase of the cell cycle by forming a complex with cyclin D (38). Palbociclib has inhibitory effects on Cushing’s disease *in vitro* (31). Clinical applications of palbociclib to Cushing’s disease are expected because cyclin D is highly expressed in aggressive PitNETs (48). We wondered if TS would show cooperative effects with palbociclib, because TS decreased the gene expressions of several cyclins, including cyclin D. In the present study, palbociclib reduced the cell viability of AtT-20 cells at lower concentrations when it was co-administered with TS than when it was not. Palbociclib is often accompanied by adverse effects such as fatigue, neutropenia, and interstitial pneumonia, which worsen as dosages increase (66, 67). TS administration may be helpful not only for tumor suppression but also for alleviating adverse effects by reducing dosages of companion drugs.

Regarding other CDK inhibitors, the therapeutic effects of seliciclib against Cushing’s disease were verified not only in a multicenter Phase 2 clinical trial, but also in an *in vitro* study (32, 33, 37). Seliciclib targets CDK2 and forms complexes with cyclin E, which is also highly expressed in ACTH-secreting PitNETs (48). A multi CDK inhibitor, flavopiridol, also demonstrated efficacy in Cushing’s disease (31). In the present study, seliciclib and flavopiridol did not exhibit cooperative effects when co-administered with TS. These results are not surprising because seliciclib and flavopiridol both induce G2/M arrest, which overlaps with the mechanism of TS (68–72).

A major limitation of the present study is that we used AtT-20, the only available ACTH-secreting murine cell line. The application of our results to clinical settings requires further investigation using materials derived from humans. Another limitation regarding the HTS results is that the ACTH reporter assay may underestimate or overestimate the levels of compounds that directly modulate cAMP. In the experiment whose results are shown in Figure 2D, we did demonstrate that the hit compounds did not directly alter cAMP production. In any case, the hit compounds may clarify aspects of the pathophysiology of ACTH-secreting PitNETs. Most of our hit compounds seemed to inhibit ACTH secretion by decreasing cell viability. Compounds that inhibited ACTH secretion while maintaining cell viability may be valuable to clarify the mechanisms of autonomous ACTH secretion by ACTH-secreting PitNET.

In conclusion, we performed HTS using an original ACTH reporter assay and identified a number of bioactive compounds that may contribute to Cushing’s disease. Of the hit compounds, we examined TS in detail and verified its therapeutic effects in *in vitro* and *in vivo* models of Cushing’s disease. In AtT-20 cells, TS induced cell cycle arrest at the G2/M phase and then apoptosis that were mediated by FOXM1-independent mechanisms including downregulation of cyclins. TS exhibited cooperative antitumor effects when co-administered with a CDK4/6 inhibitor. We believe that the list of hit compounds and the insight into the mechanisms of TS will serve as a valuable foundation for future research.

## Materials and Methods

### Plasmid construction

We cloned the coding sequences of murine *Mc2r*, *Mrap*, and *Foxm1* into a pcDNA3.1 vector (Thermo Fisher Scientific, Waltham, MA) using PCR with primers containing restriction enzyme sites and the Kozak sequence (Supplementary Table 3). These vectors were verified by sequencing. Pink Flamindo was a gift from Tetsuya Kitaguchi (Addgene plasmid #102356) (17). A large amount of these plasmid vectors was prepared using a NucleoBond Xtra Midi Plus kit (Macherey-Nagel, Düren, Germany). Concentrations of plasmid DNA were measured with a Nanodrop-1000 spectrophotometer (Thermo Fisher Scientific).

### Cells and reagents

AtT-20, a murine cell line derived from an ACTH-secreting pituitary tumor, was obtained as AtT-20/D16v-F2 from the American Type Culture Collection (Manassas, VA). GH3, a rat tumor cell line secreting somatotropin and prolactin, and MCF-7, a human breast cancer cell line, were purchased from JCRB Cell Bank (Osaka, Japan). HEK293T cells used in the present study were authenticated with short tandem repeat profiling at the JCRB Cell Bank (43, 73). These cells were maintained in Dulbecco’s modified Eagle’s medium (DMEM) (Sigma-Aldrich, St. Louis, MO) supplemented with an antibiotic-antimycotic (Thermo Fisher Scientific) and 10% fetal bovine serum (FBS) (Sigma-Aldrich) at 37°C in a humidified atmosphere containing 5% CO_2_. Recombinant ACTH was purchased from Daiichi Sankyo Co., Ltd. (Tokyo, Japan), and forskolin was procured from Tokyo Chemical Industry Co., Ltd. (Tokyo, Japan). Hoechst 33342 was purchased from Nacalai Tesque (Kyoto, Japan). Thiostrepton (Cayman Chemical, Ann Arbor, MI), palbociclib (MedChemExpress, Monmouth Junction, NJ), seliciclib (Selleck Chemicals, Houston, TX), and flavopiridol (Selleck Chemicals) were dissolved in dimethyl sulfoxide (DMSO).

### ACTH reporter assay

HEK293T cells were seeded in 100-mm dishes at 1.0 × 10^6^ cells/well in 10 mL of antibiotic-free DMEM with 10% FBS. After incubation for 24 h, transfections with 2.5 µg of pcDNA3.1-*Mc2r*, pcDNA3.1-*Mrap*, and Pink Flamindo were performed using 25 µg of PEI MAX (Polysciences, Warrington, PA). Following additional incubation for 24 h, transfected cells were harvested using phenol red-free DMEM and seeded at a density of 1.0 × 10^4^ cells/well onto CellCarrier^TM^-96 Ultra microplates (Perkin Elmer, Waltham, MA). Then, high-content non-confocal imaging was performed using the Opera Phenix High-Content Screening System (Perkin Elmer) with a 20× water immersion lens and 568-nm lasers, along with Phenix emission filters (570–630 nm) to detect the fluorescence of ACTH reporter cells. Images were acquired before and after the administration of reagents or conditioned media. Nine fields of view were imaged per well. Acquired images were analyzed using Harmony analytical software (v4.9, Perkin Elmer). Optical correction was performed using flat-field and bright-field correction. Individual cells were identified based on the Alexa 568 signal. The “find cells building block” (algorithm method M, common threshold 1.0, diameter > 30 μm, splitting sensitivity 0.2) was employed to identify cells that met the set criteria. Changes in mean fluorescence intensity of individual cells were calculated from values before and after the administration of reagents or conditioned media.

### Chemical library

The chemical library consisted of 2480 bioactive compounds obtained from Prestwick Chemical (Illkirch-Graffenstaden, France), Calbiochem (Merck Millipore, Darmstadt, Germany), and Selleck Chemicals (43). Stock solutions (10 mM) were prepared in DMSO and arrayed in 96-well plates. Dilution series were prepared by 1:10 dilution with DMSO.

### High-throughput screening

AtT-20 cells were seeded in 96-well plates at 1.0 × 10^4^ cells/well in 200 μL of phenol red-free DMEM (Nacalai Tesque) with 10% FBS. The chemical library was diluted at a 1:200 ratio with phenol red-free DMEM containing 10% FBS, and 50 μL of the diluted chemical library was added in duplicate to the 96-well plates with the seeded cells, resulting in a final concentration of 10 μM for the chemical library. DMSO was added as a negative control. The 96-well plates were then incubated for 24 h. Following incubation, we evaluated ACTH levels in conditioned media using the ACTH reporter assay described above.

### Cell viability assay

AtT-20 cells were suspended in 100 μL of DMEM supplemented with 10% FBS in 96-well plates (1.0 × 10^4^ cells/well). Cells were then treated with the reagents for 24 h or 48 h. Cell viability was evaluated at the end of treatment using Cell Count Reagent SF (Nacalai Tesque) and measured with a Spark multimode microplate reader (Tecan, Männedorf, Switzerland). This protocol was also adapted for GH3 and MCF-7 cells.

### Reagent administration and sample collection

AtT-20 cells were seeded in 12-well plates at 2.0 × 10^5^ cells/well, or in 6-well plates at 4.0 × 10^5^ cells/well. TS was added to plates with the seeded cells and the plates were then incubated for 24 h or 48 h. Nuclear staining with Hoechst 33342, RNA extraction, or protein extraction was performed after incubation. Twelve-well plates were used for RNA extraction, and six-well plates were used for nuclear staining and protein extraction.

### Measurement of ACTH and corticosterone concentrations

ACTH concentrations in conditioned media and murine plasma were analyzed using an EIA kit (Phoenix Pharmaceuticals Inc., Belmont, CA) and an ELISA kit (MD Bioproducts, Zürich, Switzerland), respectively. Plasma corticosterone concentrations were analyzed using an ELISA kit (Enzo Life Sciences, Farmingdale, NY). All procedures were performed according to the manufacturer’s protocol.

### AtT-20 xenograft mouse model

Eight-week-old KSN/Slc mice were procured from Japan SLC, Inc. (Hamamatsu, Japan) and housed in a controlled environment with a 12-hour light/dark cycle with constant humidity and temperature. KSL/Slc mice were subcutaneously injected in the right flank with 5 × 10^6^ AtT-20 cells in PBS. Mice with tumors whose volume was less than 25 mm^3^ at 1 week after transplantation were divided into two groups for experiments. One group received 100 μL of vehicle solution consisting of 20% N,N-dimethylacetamide (Sigma-Aldrich), 70% polyethylene glycol 400 (Nacalai Tesque), and 10% polysorbate 80 (Nacalai Tesque) via intraperitoneal injection every other day. The other group was administered 300 mg/kg TS dissolved in the same solution as previously reported (74). Twice a week, mice were weighed and their tumor sizes were measured using calipers. Tumor volumes were calculated using the following formula: length × width × height × 4π/3. On day 14, between 10:00 and 13:00, mice were sacrificed by isoflurane exposure. To obtain plasma, blood samples were collected in microtubes containing aprotinin (Sigma-Aldrich) and ethylenediaminetetraacetic acid (Thermo Fisher Scientific), left undisturbed for 30 min at room temperature and then on ice. After centrifugation at 3,000 *g* for 15 min at 4°C, the supernatants were collected and stored at −80°C until use. The pituitary glands were harvested for RNA extraction, immediately frozen in liquid nitrogen, and stored at −80°C until needed. Every effort was made to minimize suffering. All experimental procedures involving animals were approved by the Animal Research Committee, Kyoto University Graduate School of Medicine (permit number: MedKyo07598). Care of animals and all animal experiments were conducted in accordance with our institutional guidelines.

### Quantitative RT-PCR

Total RNA was extracted using a Nucleospin RNA Plus kit (Macherey-Nagel) and reverse transcribed using ReverTra Ace (TOYOBO Life Science, Osaka, Japan). Quantitative RT-PCR was performed using THUNDERBIRD SYBR qPCR MIX (TOYOBO Life Science) with the StepOnePlus Real-time PCR System (Thermo Fisher Scientific). Results were normalized using the means of *Actb* and *Gapdh* as reference genes. Relative mRNA expression was evaluated using the comparative threshold cycle method. The primers used are listed in Supplementary Table 3.

### Western blotting

AtT-20 cells were lysed in radioimmunoprecipitation assay buffer (Nacalai Tesque) and left on ice for 30 min. Supernatants were centrifuged at 10,000 *g* for 10 min at 4°C and collected in microtubes. Protein concentrations were assessed using the Protein Assay BCA kit (Nacalai Tesque) with bovine serum albumin (Bio-Rad, Hercules, CA) as a standard. For FOXM1, 50 μg/lane of lysates were electrophoresed on NuPAGE 3–8% Tris-Acetate Gels (Thermo Fisher Scientific), while 10 μg/lane of lysates were electrophoresed on Bolt 4–12% Bis-Tris Plus Gels (Thermo Fisher Scientific) for β-actin. The protein samples were then transferred onto polyvinylidene difluoride membranes using the iBlot2 Dry Blotting System (Thermo Fisher Scientific). After blocking the membranes with Blocking One (Nakalai Tesque) for 30 min at room temperature, they were incubated overnight at 4°C with primary antibodies. The primary antibodies used were a mouse monoclonal anti-FOXM1 antibody (1:500, sc-376471; Santa Cruz Biotechnology, Dallas, TX), and a mouse monoclonal antibody anti–β-actin antibody (1:2000, 010-27841; FUJIFILM Wako Pure Chemical, Osaka, Japan) as an endogenous control. Subsequently, membranes were incubated with a horseradish peroxidase–conjugated secondary antibody (1:2000, sc-5254409, Santa Cruz Biotechnology) for 2 h at room temperature. Can Get Signal (TOYOBO Life Science) was used for detection of FOXM1 bands. Bands were detected using a chemiluminescent method with Chemi Lumi One Super (Nacalai Tesque) in ImageQuant LAS 4000 (GE Healthcare, Chicago, IL).

As a positive control for FOXM1, we prepared AtT-20 cells transfected with pcDNA3.1-*Foxm1*. In detail, AtT-20 cells were seeded in 100-mm dishes at 1.0 × 10^6^ cells/dish in 10 mL of antibiotic-free DMEM supplemented with 10% FBS. After incubation for 24 h, transient transfections of 5 μg of pcDNA3.1-*Foxm1* were performed using 25 μg of Lipofectamine 2000 (Thermo Fisher Scientific). After 24 h, proteins were collected by the method described above.

### siRNA transfection

An aliquot of 5 pmol siRNA specific for murine *Foxm1* (#4390771) (Thermo Fisher Scientific) or a negative control siRNA (#4390846) (Thermo Fisher Scientific) was transfected into AtT-20 cells (2.0× 10^5^ cells/well) using 12-well plates and 3 μL of the Lipofectamine RNAiMAX reagent (Thermo Fisher Scientific) by reverse transfection. To determine the efficacy of knockdown by siRNA, RNA was extracted 48 h after siRNA transfection and analyzed by quantitative RT-PCR as described above. Changes in protein levels of FOXM1 following siRNA transfections were evaluated by western blot.

### RNA-seq and bioinformatics analysis

mRNA paired-end libraries were constructed using the TruSeq Stranded mRNA Sample Prep Kit (Illumina, Inc., San Diego, CA). Paired-end sequencing of 100 bp was performed on a NovaSeq 6000 system (Illumina, Inc.). Quality control metrics of raw sequencing reads were performed with FastQC v0.11.7, and low-quality reads were removed with Trimmomatic 0.38. Then, reads were mapped to the mm10 reference sequence with HISAT2 version 2.1.0 and were assembled into transcripts with StringTie version 2.1.3b. The expression profile was calculated as transcript per kilobase million. For DEGs analyses, values of fold changes were calculated and then the DEGs with an adjusted p < 0.05 and fold changes ≤ −2 or ≥ 2 were determined using DESeq2. Gene-set enrichment analyses were performed based on the KEGG database and GO with gProfiler. These experiments and analyses were conducted by Macrogen Japan (Tokyo, Japan).

### Flow cytometry

AtT-20 cells were plated in six-well plates at a concentration of 4.0 × 10^5^ cells/well and treated with several concentrations of TS or palbociclib for 48 h. For cell cycle analysis, samples were stained with propidium iodide (Nacalai Tesque) for 30 min at 4°C after ethanol fixation and digestion by Ribonuclease A Solution (Nacalai Tesque). The Annexin V-FITC Apoptosis Detection Kit (Nacalai Tesque) was used for the apoptosis assay. The cell cycle phase distribution and apoptosis were detected with an SH800 cell sorter (Sony Biotechnology Inc., Tokyo, Japan), and analysis was performed using the FlowJo 10.7.1 software (BD Biosciences, Franklin Lakes, NJ).

### Statistical analysis

All results are expressed as the mean ± SEM. Statistical analysis of the data was performed using the Student’s *t*-test, ANOVA followed by the Tukey-Kramer test, or the Dunnett test. JMP Pro version 17.0.0 (SAS Institute Inc., Cary, NC) was used for statistical analyses. Statistical analyses of DEGs were performed with the Wald test using DESeq2. Fisher’s exact test was used for enrichment analyses. Statistical significance was defined as *p* < 0.05.

## Acknowledgments

HTS was performed with the support of Yukiko Okuno at the Medical Research Support Center, Kyoto University Graduate School of Medicine. HTS was partially supported by the Platform Project for Supporting Drug Discovery and Life Science Research, Basis for Supporting Innovative Drug Discovery and Life Science Research (BINDS), from AMED under grant no. JP23ama121034 (support no. 2292). Grants from the Japan Foundation for Applied Enzymology supported this work.

## Author Contributions

TH conducted the experiments and analyzed the data. IY designed the experiments and provided funding. DK, TS, HF, KO, YU, TF, and DT contributed to the discussion. NI supervised the entire study. TH and IY drafted the manuscript, and all authors reviewed, edited, and approved the manuscript.

## Disclosures

The authors have nothing to disclose.

## Data Availability

Some or all data sets generated and/or analyzed during the current study are not publicly available but are available from the corresponding author on reasonable request.

## Notes

### Competing Interest Statement

The authors have declared no competing interest.

