## Supplementary Figures and Table 3 for "High-throughput screening for Cushing’s disease: therapeutic potential of thiostrepton via cell cycle regulation"

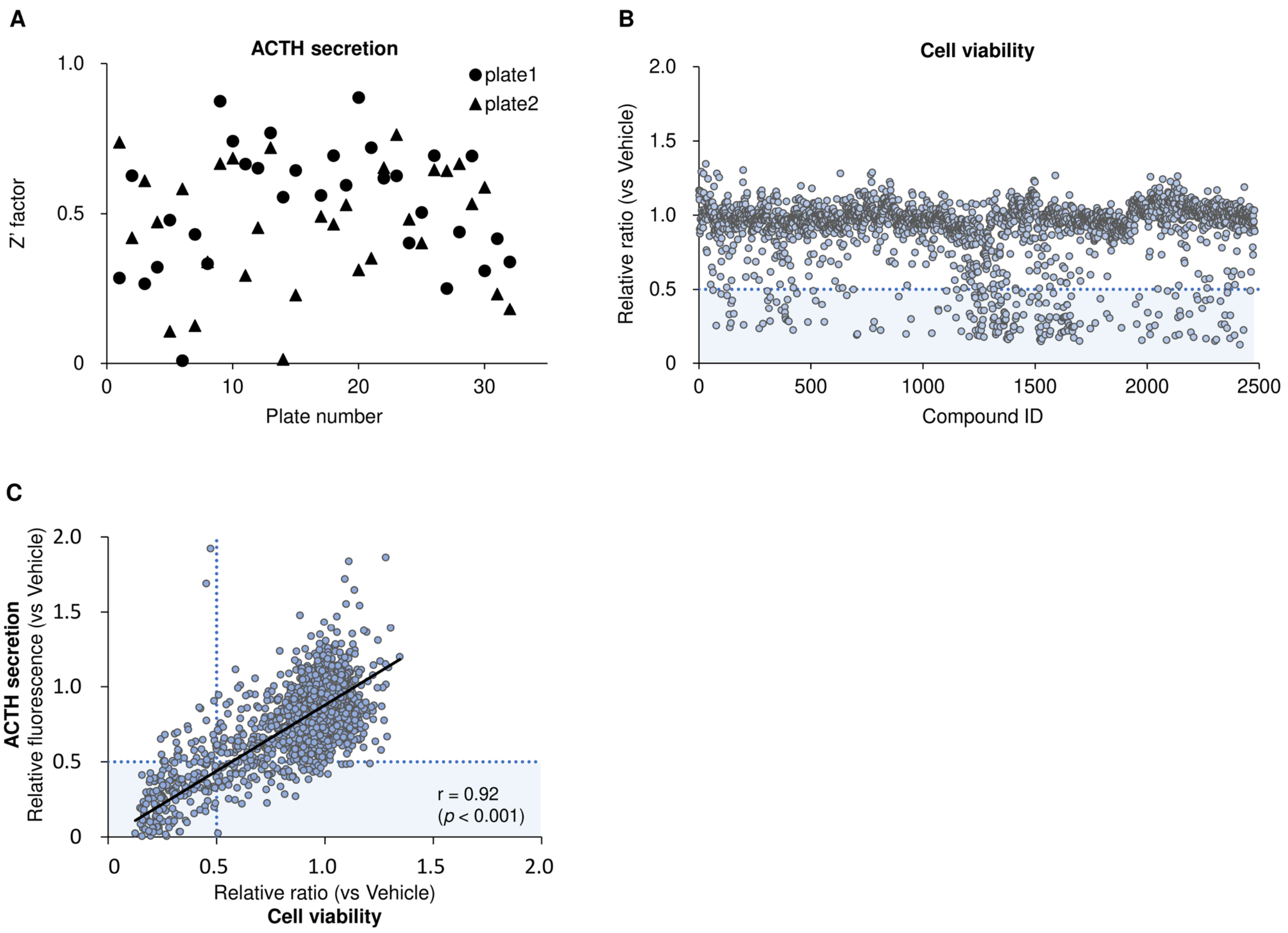


**Supplementary Figure 1.** (A) Z' factor in the first screening of high throughput screening (HTS). Compounds were added to AtT-20 cells seeded in 96-well plates in duplicate (Plates 1 and 2) for each plate of the chemical library. (B) Cell viability of AtT-20 cells treated with all 2480 compounds. Values were normalized as relative ratio; in each plate, 1.0 was the result of AtT-20 cells treated with vehicle. (C) A scatter plot of HTS results. A vertical axis represents changes in ACTH secretion determined by the ACTH reporter assay and a horizontal axis represents cell viability. Values of ACTH secretion were normalized as relative fluorescence; in each plate, 1.0 was the result of AtT-20 cells treated with vehicle. Correlation was analyzed by calculating Pearson correlation coefficient.

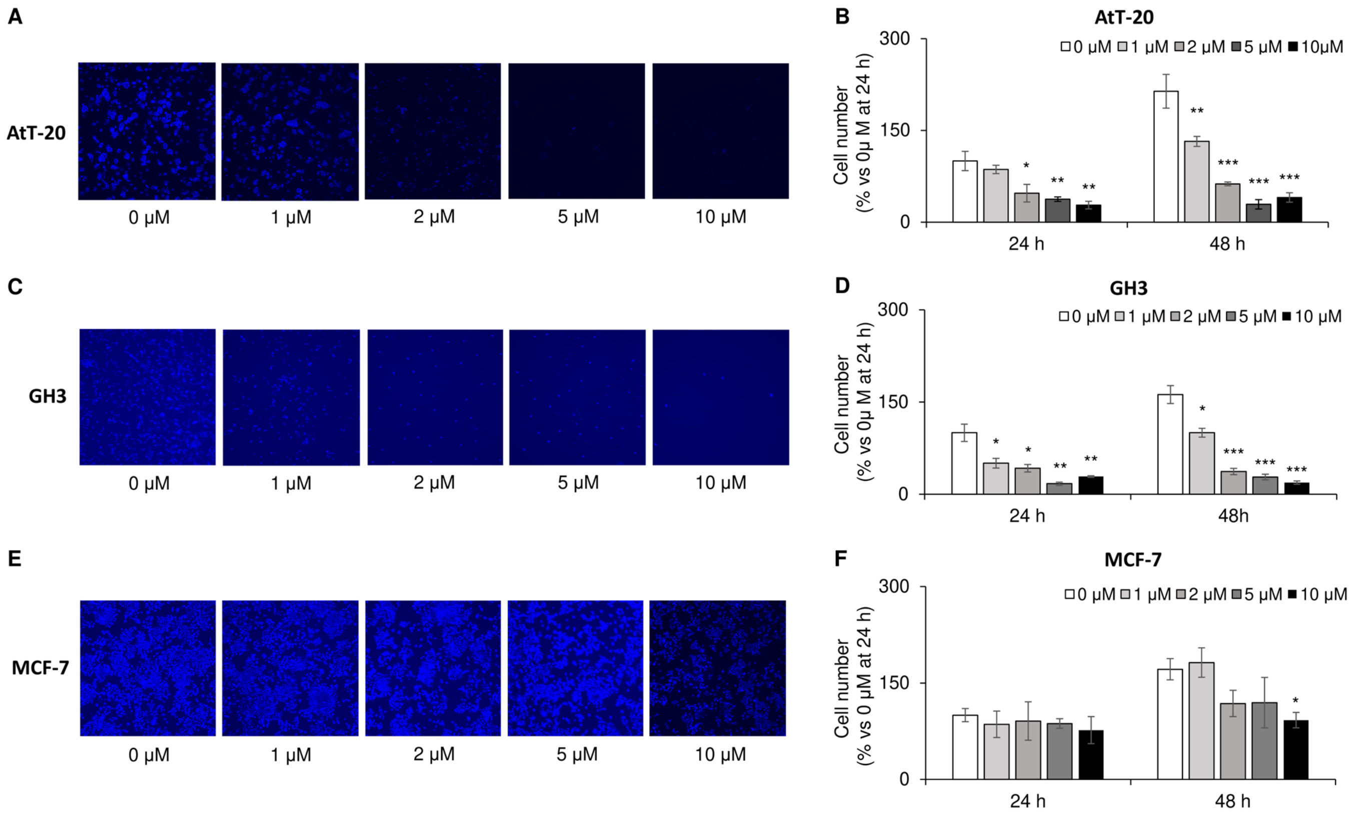


**Supplementary Figure 2.** (A) Nuclear staining with Hoechst 33342 of AtT-20 cells treated with a dilution series of thiostrepton (TS) after 48 h. (B) Numbers of AtT-20 cells treated with TS were counted after 24 and 48 h. Results are shown as the ratio relative to 0 µM at 24 h. $n = 4$ each. (C, D) Nuclear staining with Hoechst 33342 (C) and numbers (D) of GH3 cells treated with TS. $n = 4$ each. (E, F) Nuclear staining with Hoechst 33342 (E) and numbers (F) of MCF-7 cells treated with TS. $n = 4$ each. Data are represented as means ± SEM. Statistical analyses were performed using ANOVA followed by the Dunnett test with comparisons to results without TS administration. The $p$-value is presented as $*p < 0.05$, $**p < 0.01$, and $***p < 0.001$.

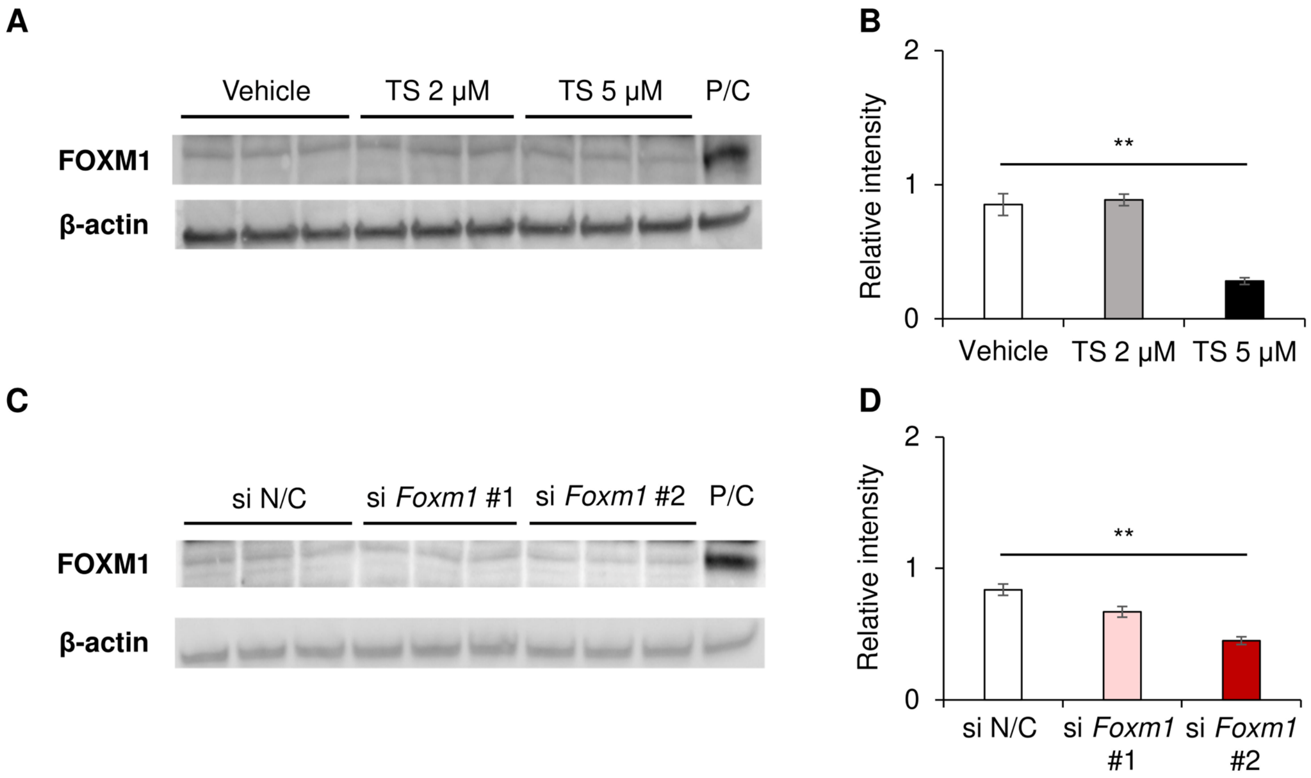


**Supplementary Figure 3.** (A, B) Western blot analyses of AtT-20 cells treated with TS. A positive control sample (P/C) was prepared by AtT-20 cells transfected with pcDNA3.1-*Foxm1*. FOXM1 expressions were evaluated using values normalized to β-actin expressions. (C, D) Western blot analyses of AtT-20 cells with *Foxm1* knockdown by transfection with short interfering RNA (siRNA). We used a negative control siRNA (si N/C) and two siRNAs for *Foxm1* (si *Foxm1* #1 and si *Foxm1* #2). Protein samples were collected after 24 h. Data are represented as means ± SEM. Statistical analyses were performed using ANOVA followed by the Tukey-Kramer test. The $p$-value is presented as $^{**}p < 0.01$.

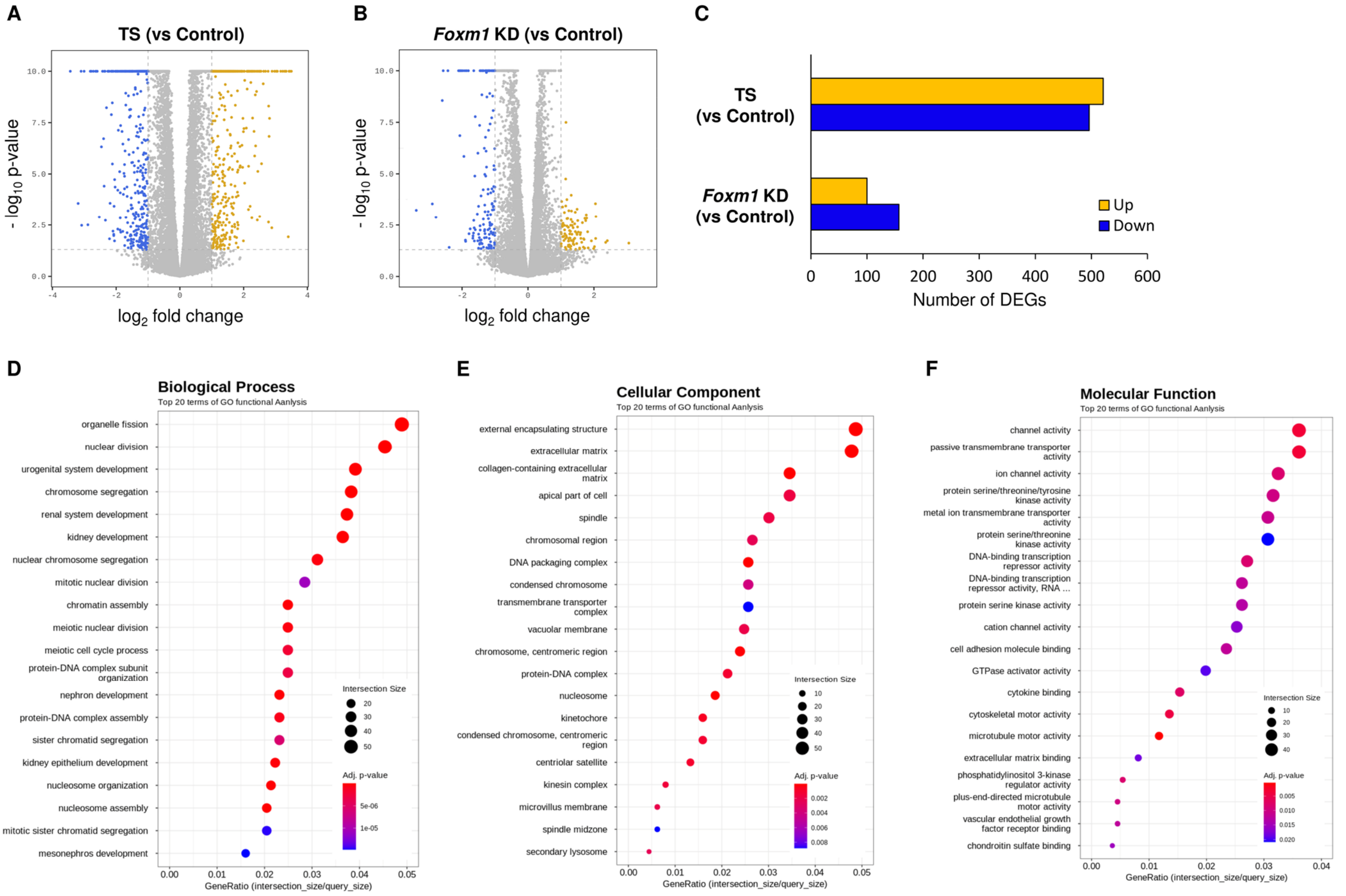


**Supplementary Figure 4.** (A–C) Differentially expressed genes (DEGs) determined in RNA-sequencing (RNA-seq) analyses. Volcano plots of DEGs of the TS group treated with 2 μM TS (A) and the *Foxm1* knockdown (KD) group transfected with siRNA for *Foxm1* (B), and numbers of DEGs (C). (D–F) The top 20 terms in enrichment analyses using gene ontology (GO). DEGs were determined by comparisons with the Control group without either TS administration or *Foxm1* knockdown. Statistical analyses were performed using the Wald test.

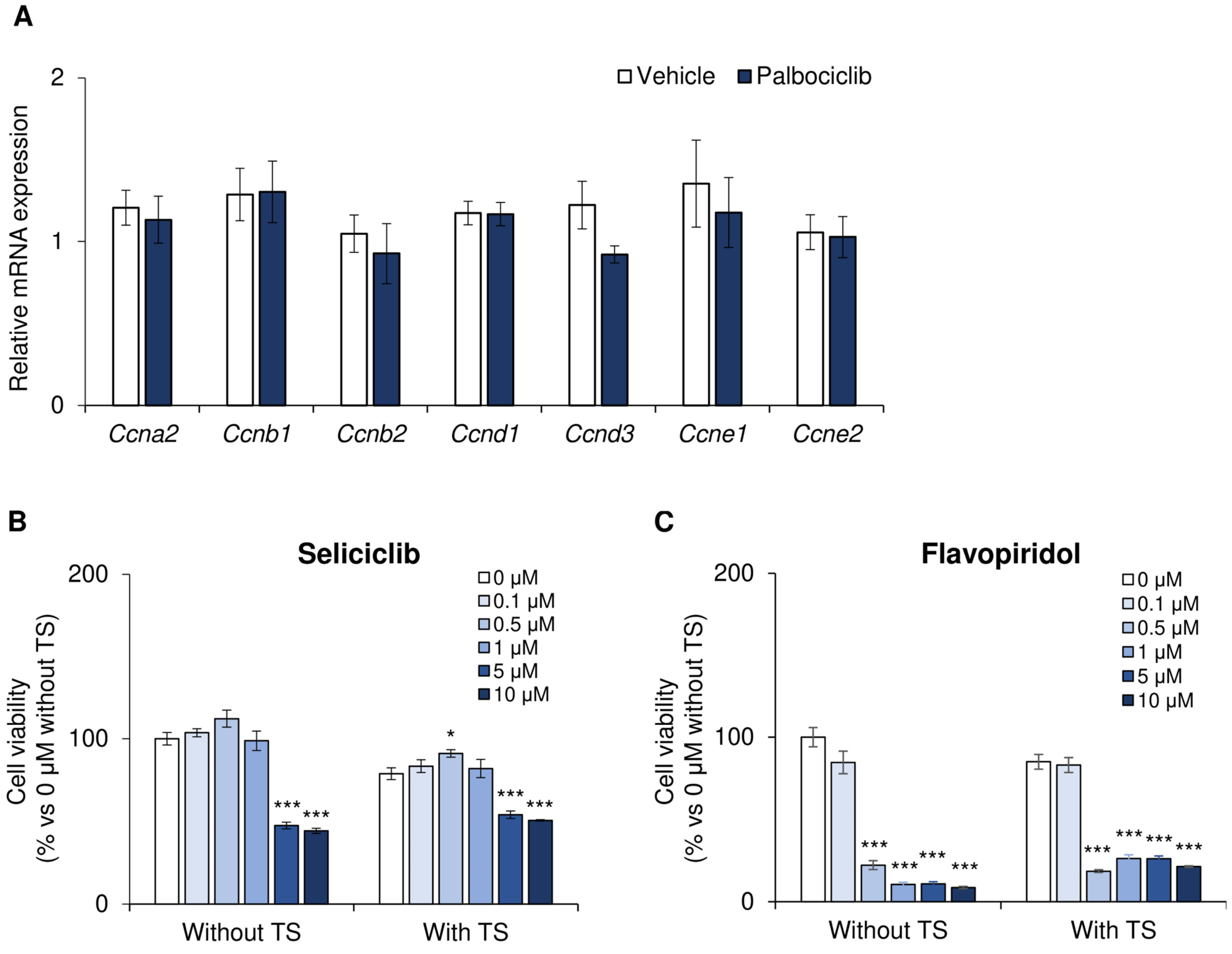


**Supplementary Figure 5.** (A) Quantitative RT-PCR of mRNAs of cyclins in AtT-20 cells treated with 5 μM palbociclib for 24 h. Data are normalized by means of *Actb* and *Gapdh* and shown as the ratio relative to vehicle. $n = 3$ each. (B, C) Cell viability of treated AtT-20 cells. Either vehicle or 2 μM TS was co-administered with a dilution series of seliciclib (B) and flavopiridol (C) for 48 h. Results are shown as the ratio relative to 0 μM without TS. $n = 6$ each. Data are represented as means ± SEM. Statistical analyses were performed using the Student's *t*-test for a panel A and ANOVA followed by the Dunnett test with comparisons to results without administration of seliciclib or flavopiridol for panels B and C. The *p*-value is presented as $*p < 0.05$ and $***p < 0.001$.

**Supplementary Table 3. Primers used for plasmid construction and quantitative RT-PCR**

| Genes (accession number) | |
|---|---|
| Plasmid construction | |
| *Mc2r* (NM_008560) | Fw: AGGAGGATCCGCCACCATGAAGCATATTATCAAT<br>Rv: AGGAGAATTCCTAATACCGGTTGCAGAA |
| *Mrap* (NM_029844) | Fw: AATAGGATCCGCCACCATGGCCAACGGGACCGACGC<br>Rv: AATAGAATTCCTAGGGGAGAGCCAGGGGC |
| *Foxm1* (NM_008021) | Fw: AATAGCTAGCGCCACCATGAGAACCAGCCCCCGCCG<br>Rv: AATAGGATCCCTAAGGGATGAACTGAGACCA |
| Quantitative RT-PCR | |
| *Foxm1* (NM_008021) | Fw: CACTTGGATTGAGGACCACTT<br>Rv: GTCGTTTCTGCTGTGATTCC |
| *Pomc* (NM_008895) | Fw: TGTACCCCAACGTTGCTGAG<br>Rv: AGGACCTGCTCCAAGCCTAA |
| *Gh* (NM_008117) | Fw: GCTTCTCAGAGACCATCCCG<br>Rv: ATCCTGCTGAGGAACTGCAC |
| *Lhb* (NM_008497) | Fw: TCTGCATCACCTTCACCACC<br>Rv: CTGAGGCACAGGAGGCAAA |
| *Fshb* (NM_008045) | Fw: ATACCACTTGGTGTGCGGG<br>Rv: TACTTTCTGGGTATTGGGCCG |
| *Prl* (NM_011164) | Fw: AGAGCTGTTTGACCGTGTGG<br>Rv: GTAGCCAGGGAAGAAGTGGG |
| *Tshb* (NM_009432) | Fw: ACAGACATCCTGAGAGAGTGC<br>Rv: GAGAGTGTGCCTACTGCCTG |
| *Pitx1* (NM_011097) | Fw: GAGAACTCCGCCAGCGAATC<br>Rv: CTCTGGCCCCTTGGCTTC |
| *Tbx19* (NM_032005) | Fw: CCTTTCGCCAAGGCCTTCTT<br>Rv: AGTAGGTCACATGCTGGCTC |
| *Crhr1* (NM_007762) | Fw: GGGCAGCCCGTGTGAATTAT<br>Rv: GAAGAGGACAAAGGCCACCA |
| *Ccna2* (NM_009828) | Fw: GGGTTCTTCTCTGGCTCCAA<br>Rv: AAAGAGTGTCAGCCTCCGGG |
| *Ccnb1* (NM_172301) | Fw: TCTTGACAACGGTGAATGGA<br>Rv: CTTTGTGAGGCCACAGTTCA |
| *Ccnb2* (NM_007630) | Fw: AGCTCTGCAGTGACTACGTG<br>Rv: CTGCAGAAGCCGAAACTTGG |
| *Ccnd1* (NM_007631) | Fw: CAGATTGAAGCCCTTCTGGA<br>Rv: ACCAGCCTCTTCCTCCACTT |
| *Ccnd3* (NM_007632) | Fw: GCGTGCAAAAGGAGATCAAGCC<br>Rv: CCAGGTAGTTCATAGCCAGAGG |

| | |
|---|---|
| *Ccne1* (NM_007633) | Fw: AAATCAGACCACCCAGAGCCT<br>Rv: TGGAGCTTATAGACTTCGCACACC |
| *Ccne2* (NM_009830) | Fw: TCTGTGCTGCCAGCTGTAAT<br>Rv: TGTCCCTCCAGGTAACCATACT |
| *Actb* (NM_007393) | Fw: CATCCGTAAAGACCTCTATGCCAAC<br>Rv: ATGGAGCCACCGATCCACA |
| *Gapdh* (NM_008084) | Fw: TGCACCACCAACTGCTTAGC<br>Rv: GGATGCAGGGATGATGTTCTG |

Fw; Forward primer (5′–3′), Rv; Reverse primer (5′–3′).
